# Evaluation of Whole Exome Sequencing as an Alternative of BeadChip and Whole Genome Sequencing in Human Population Genetic Analysis

**DOI:** 10.1101/280552

**Authors:** Zoltán Maróti, Zsolt Boldogkői, Dóra Tombácz, Michael Snyder, Tibor Kalmár

## Abstract

Understanding the underlying genetic structure of human populations is of fundamental interest to both biological and social sciences. Advances in high-throughput genotyping technology have markedly improved our understanding of global patterns of human genetic variation. The most widely used methods for collecting variant information at the DNA-level include whole genome sequencing, which continues to remain costly, and the more economical solution of array-based techniques, as these are capable of simultaneously genotyping a pre-selected set of variable DNA sites in the human genome. The largest publicly accessible set of human genomic sequence data available today originates from exome sequencing that comprises around 1.2% of the whole genome (approximately 30 million base pairs). In this study, we compared the application of the exome dataset to the array-based dataset and to the gold standard whole genome dataset using the same population genetic analysis methods. Our results draw attention to some of the inherent problems that arise from using pre-selected SNP sets for population genetic analysis. Additionally, we demonstrate that exome sequencing provides a better alternative to the array-based methods for population genetic analysis. In this study, we propose a strategy for unbiased variant collection from exome data and offer a bioinformatics protocol for proper data processing.

## INTRODUCTION

The investigation of the ethnogenesis of human populations is made possible by population genetic studies, through comparing genetic makeup and frequencies of the selected variants or alleles, and also by computing their genetic distance from the rest of the studied population or their level of admixture (Li et al. 2008; Hellenthal et al. 2014). Compared to their more costly whole genome sequencing (WGS) counterparts, these assays predominantly use array-based genotyping techniques (various Human BeadChip arrays /610, 640, 650, 1M/, Infinium Multi-Ethnic BeadChip arrays from Illumina, Affymetrix Genome-Wide Human SNP Array, Affymetrix Human Origin Array, etc.) that include single nucleotide polymorphism (SNP) sets based on the evaluation of previous genome sequencing data (CEPH, HGP, HapMap databases), with the emphasis on population-specific, ancestry-informative markers (AIMs). AIMs were first introduced in 2008 by Halder and colleagues (Halder et al. 2008), as a panel of 176 autosomal AIMs that were capable of effectively distinguishing individual biogeographical ancestry and admixture proportions from among four continental ancestral populations. The importance of AIMs has palpable significance in the medical field as well. While case-control design studies can be an efficacious strategy for identifying candidate genes in complex diseases in a population, in diversely admixed populations (e.g. Latin Americans, with admixture of American Indians, Europeans and Africans) population stratification can affect association studies and thereby could lead to false genetic associations (Stefflova et al. 2011). This undesirable distortion can be minimized by genotyping AIMs.

Ultimately, application of the Whole Exome Sequencing (WES) method had spread and gained popularity, as WES is cost effective for routine genetic diagnosis of rare inherited diseases, and extensive databases have been generated containing thousands of publicly accessible exomes /Exome Aggregation Consortium ~61000 exomes (Lek et al. 2016), Exome Variant Server ~6500 exomes (Exome Variant Server)/. A typical WES data contains 100,000 to 130,000 variants (if all the high coverage reads are also analyzed, including the flanking regions of exons). In rare disease diagnostics, focus is on filtering out all non-disease specific variants and to find the one or two causative mutations that lead to the disease phenotype. In practice, all rare and common variants aid the exploration of disease variants only as population controls. Since exome data by definition also contains all of the functional variants that are under selection pressure, in this study, we explored whether this could lead to any bias in population genetic analysis.

Thus, we assessed the usability of exome data in comparison with the other approaches in population genetic analyses. In order to compare the practicality of each strategy – (the genome data as unbiased standard, the commonly used array data, and the potentially usable exome data) – we generated three different types of datasets, all based on the same publicly available experimental data (HGP ~2,500 unrelated genomes). The results we obtained from these datasets were compared using the most widely used population genetic calculations (including Fixation index (F_ST_), Principal Components Analysis (PCA), *f*_*3*_- and *f*_*4*_-statistics, admixture) by utilizing the commonly used tools: EIGENSOFT (Patterson et al. 2006), ADMIXTOOLS (Patterson et al. 2012), Admixture (Alexander et al. 2009) and TreeMix (Pickrell and Pritchard 2012).

Migration, admixture, adaptation, and genetic drift lead to genetic diversity between human populations. By studying genetic diversity within and between populations, we could reconstruct how these populations are related to one another. Fixation index a measure of genetic structure developed by Sewall Wright (Wright 1951; Slatkin 1991) can be estimated from genetic polymorphism data. F_ST_ is the proportion of the total genetic variance contained in a subpopulation (S) relative to the total genetic variance (*T*). Its values can range from 0 to 1, where F_ST_ = 0 implies panmixia (absence of any differentiation among subpopulations) and F_ST_ = 1 implies complete divergence between populations.

Principal Components Analysis is the most widely used approach for identifying ancestry differences among a group of individuals (Price et al. 2006; Novembre and Stephens 2008). When applied to genotype data, it calculates principal components (or eigenvectors), which can be viewed as continuous axes of variation that reflect genetic variation due to ancestry in the sample. Individuals with similar values for a particular top principal component will have similar ancestry for that axes. Application of principal components to genetic data from European samples (Novembre and Stephens 2008) showed that among Europeans for whom all four grandparents originated in the same country, the first two principal components computed using 200,000 SNPs could geographically map their country of origin quite accurately.

F-statistics measure shared genetic drift among sets of populations and can be used to test simple and complex hypotheses about admixture events between populations (Peter 2016). The *f*_*3*_-statistics is used for testing the relationship of three populations (Reich et al. 2009) by allowing the detection of the presence of admixture in a population C from two other populations, A and B. If the value F_3_(C; A, B) is negative, then C does not appear to form a simple tree with A and B, but instead appears to be a mixture of A and B. If F_3_ is zero, it indicates the absence of admixture, while a positive F_3_ value implies simple tree-like relations among A, B and C (however, it does not reject admixture). Because of the complex genetic ancestry of human populations, there usually exists more than one possible model for any studied case. The *f*_*4*_-statistics has been developed in order to test alternative hypothetical trees (Keinan et al. 2007). The *f*_*4*_-statistics is used to estimate the admixture proportion of a test population (*PC*) under the assumption that we have a correct historical model. In this test, *P*A and *P*B are the potential contributors and *P*I is a reference population with no direct contribution to *PC*.

Admixture analysis is based on the maximum likelihood estimation of individual ancestries from multi-locus SNP genotype datasets (Pritchard et al. 2000; Alexander et al. 2009). It estimates the best possible sources and proportions of admixing components for any hypothetical number (K) of admixing sources. Visualization of admixture components offers an insight into the genetic structure of the studied populations.

Because there is no publicly available WGS data from modern Hungarians and since our results indicate that the use of exome data is suitable for population genetic analysis, we carried out population genetic analysis of modern Hungarians based on their exome data. Based on our results, we also evaluated a strategy to filter exome data and offer the most suitable parameters for using them in population genetic analysis.

## RESULTS

### The BEADCHIP dataset contained markedly higher amount of linked markers

Some population genetic approaches like PCA or admixture analysis are based on the assumption of linkage equilibrium. Thus, it is important that linked markers are pruned from the datasets as linkage disequilibrium (LD) between tightly linked markers causes certain haplotypes to be more frequent than expected and large blocks of markers in complete LD can seriously distort the eigenvector/eigenvalue structure (Patterson et al. 2006). This is especially important in the exome dataset as exons of genes, or genes could be tightly packed in small, transcriptionally active chromosome regions. Therefore, we altered the recommended 50kb sliding window (Alexander et al. 2009) in the *–indep-pairwise* algorithm of Plink to 10,000kb, while maintaining the recommended 10 SNPs increment and r^2^ threshold of 0.1. The variant counts and the effect of LD pruning on the different datasets are summarized in Table 1.

**Table 1:**
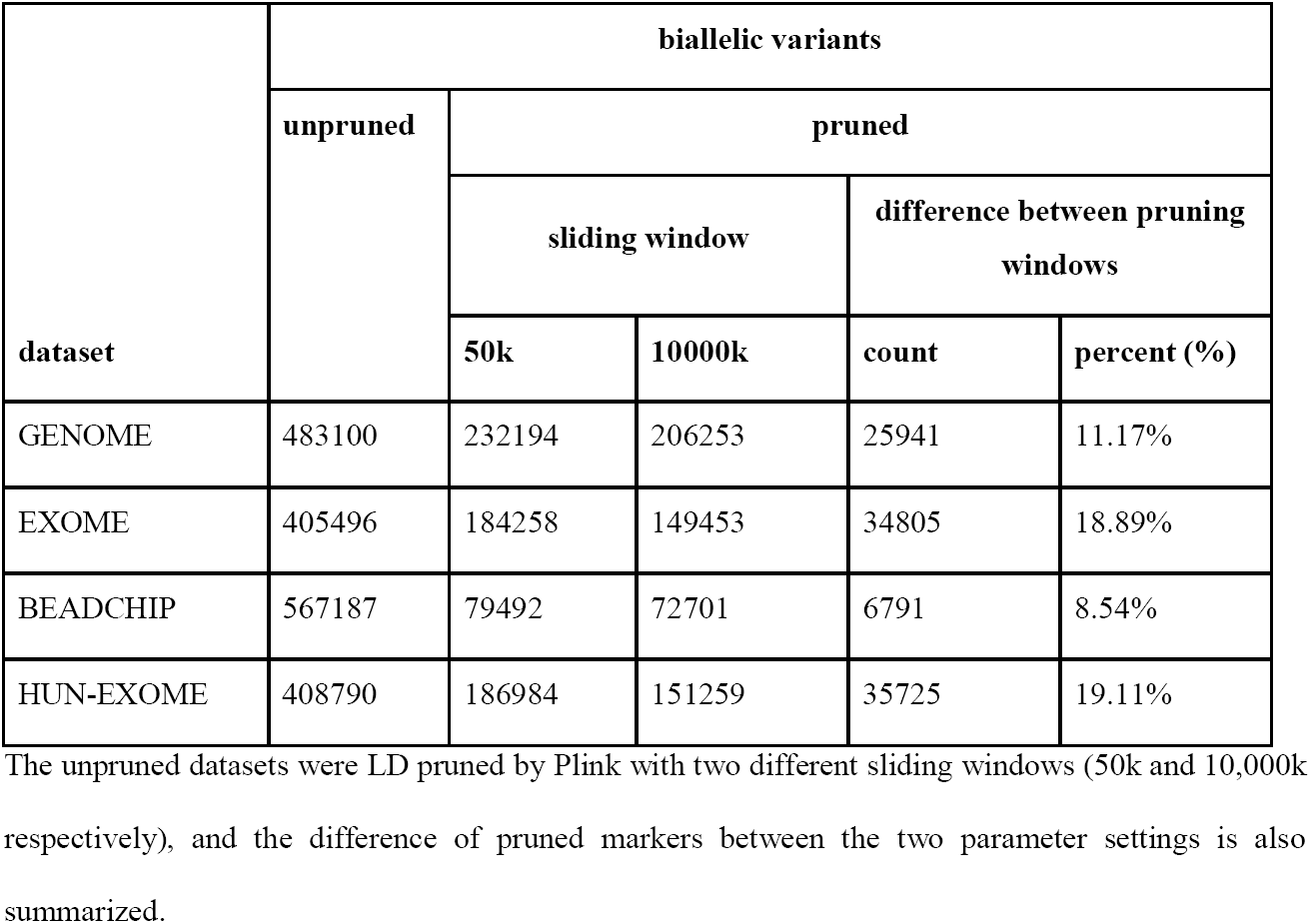
Number of high-quality biallelic variants in different datasets before and after LD pruning.

The unpruned datasets were LD pruned by Plink with two different sliding windows (50k and 10,000k respectively), and the difference of pruned markers between the two parameter settings is also summarized.

Interestingly, the BEADCHIP dataset, even with the 50kb sliding window, contained a much higher proportion of linked markers (~86%) compared to the GENOME (~52%) and EXOME (~55%) datasets. As expected, the larger pruning window affected mostly the EXOME dataset (~19% additional markers pruned), while the GENOME (~11%) and BEADCHIP (~9%) datasets were affected to a lesser degree.

### *F*_*ST*_ values based on the BEADCHIP dataset are systematically overestimated between populations with larger genetic distance

For each dataset, we calculated the pairwise F_ST_ value between each studied population and compared the results of the different datasets (Supplementary Table 1). In general, the F_ST_ distances generated from the GENOME and EXOME datasets were found to be nearly identical. However in the EXOME dataset we found very small but systematic differences between the F_ST_ values of African (except in the LWK African population) and European populations and the African and East Asian populations. We observed that F_ST_ values calculated on the basis of the BEADCHIP dataset were systematically overestimated between populations originating from different super-populations.

### Eigenvalues are notably larger for the BEADCHIP dataset compared to GENOME and EXOME datasets in PCA

For each dataset, we performed the PCA analysis of all samples without outlier removal. The different datasets show a remarkably similar overall picture by the first two eigenvectors (Figure 1A-C). The relative positions of the super-populations are almost the same, and we can even pinpoint several outlier individuals – for example, some individuals with African ancestry, marked by red dots in the middle – that are positioned in a very similar pattern in each dataset. The greatest difference is that the absolute values of eigenvectors are significantly larger for the BEADCHIP dataset compared to the GENOME and EXOME datasets, while the EXOME dataset has the most similar eigenvector values to the GENOME dataset. The complete logs of all PCA analyses on the different datasets (containing detailed Tracy-Widom statistics, average divergence between populations, ANOVA statistics for population differences along each eigenvector, statistical significance of differences between populations and list of eigen best SNPs) is summarized in Supplementary DATA/PCA. According to the detailed statistic included in the logs the three datasets showed similar statistic power to differentiate populations.

**Figure 1:**
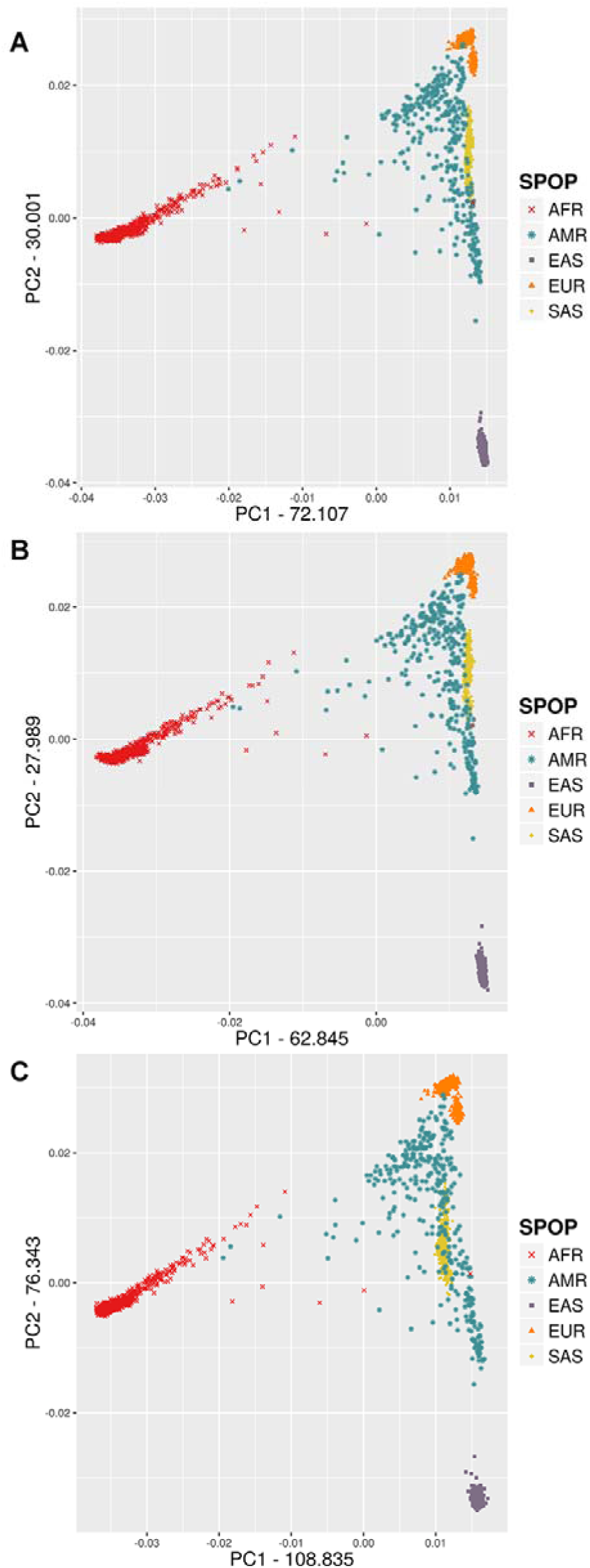
PCA analysis of super-populations using different datasets **A)** GENOME **B)** EXOME **C)** BEADCHIP.

In order to investigate whether the BEADCHIP and EXOME datasets represent similar population relations within each super-population in comparison to the GENOME dataset, next we performed the PCA analysis restricted to each super-population using default outlier removal (removing individuals with > 6 SD). The detailed PCA of the populations for each super-population showed that the indicated eigenvectors and values were very similar in all three datasets (Figure 2A-E). The greatest difference between the three datasets was seen in the AFR super-population. The differences between the overlap of the historically known admixed ASW and ACB African populations and their relations to the other African populations indicated slightly different population affiliations in the BEADCHIP dataset compared to the GENOME and EXOME datasets.

**Figure 2:**
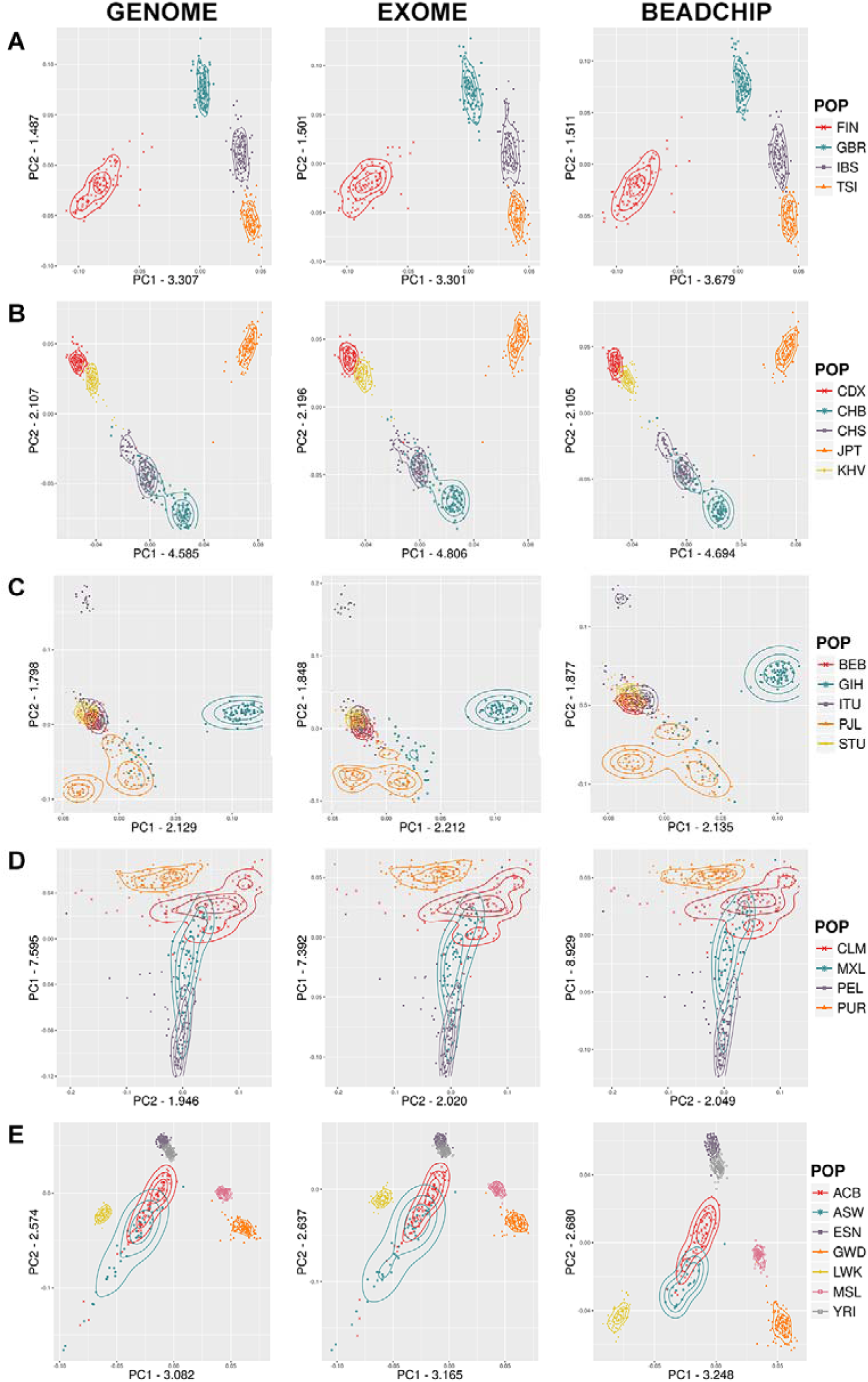
Detailed PCA of super-populations: **A)** European, **B)** East Asian, **C)** South Asian, **D)** Admixed American, **E)** African.

### *f*_3_-statistics of BEADCHIP dataset deviates from GENOME dataset

In order to compare the usability of the three datasets we calculated the *f*_*3*_-statistics for all possible combinations of population triads and plotted the resulting F_3_ values Figure 3.

**Figure 3:**
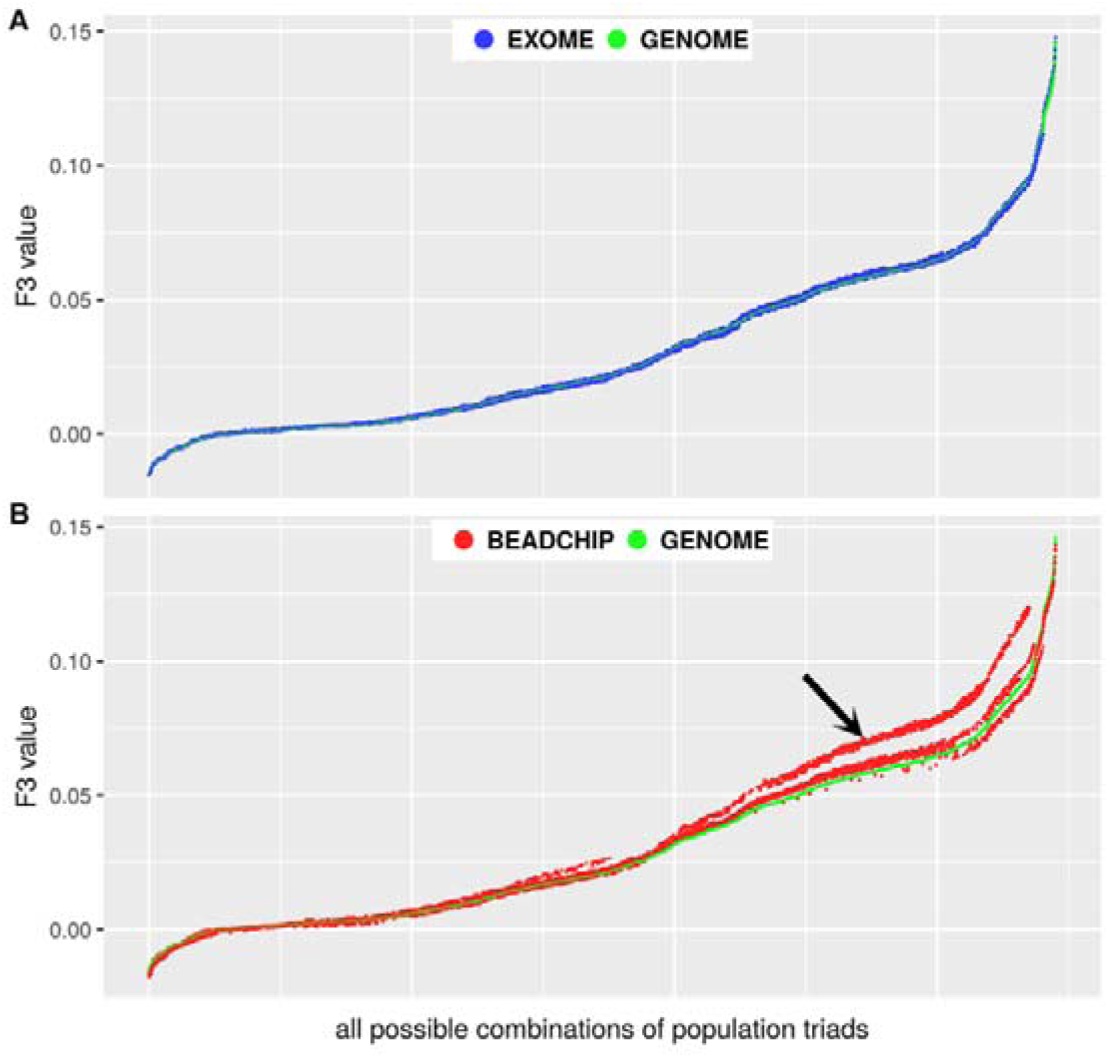
Comparison of F_3_ values obtained from the GENOME, EXOME and BEADCHIP datasets. F_3_ values were ordered and plotted relative to all possible combinations of population triads. **A)** F_3_ values from the EXOME vs. the GENOME dataset **B)** F_3_ values from the BEADCHIP vs. the GENOME dataset. The arrow denotes substantial deviations from the GENOME F_3_ values.

Figure 3A shows that the EXOME F_3_ values is almost identical to GENOME results (Pearson correlation r=0.9998), while the BEADCHIP data (Figure 3B) presents less correlation (r=0.9911) with the F_3_ values calculated from the GENOME dataset. The differences are confined to the larger positive F_3_ values in our analyzed populations (shown as deviating red dots from green dots in Figure 3B). The most deviating cases were those where F_3_ value were calculated between any two East Asian populations in relation to an arbitrary African population.

### *F*_4_ (TSI, X; CHB; YRI) values were comparable in all analyzed datasets

In the analyzed populations, all possible combinations of any four populations result in an exceedingly large in number. In most cases, the relations between these population combinations would be meaningless. Therefore, in order to test potential bias between the different datasets, we only calculated the *f*_*4*_-statistic corresponding to a well-known East Eurasian-like ancestry of Northern European populations (Reich et al. 2012). The F_4_(TSI, X; CHB; YRI) values where X denotes all possible test populations were calculated for each dataset (Figure 4 A-C).

**Figure 4:**
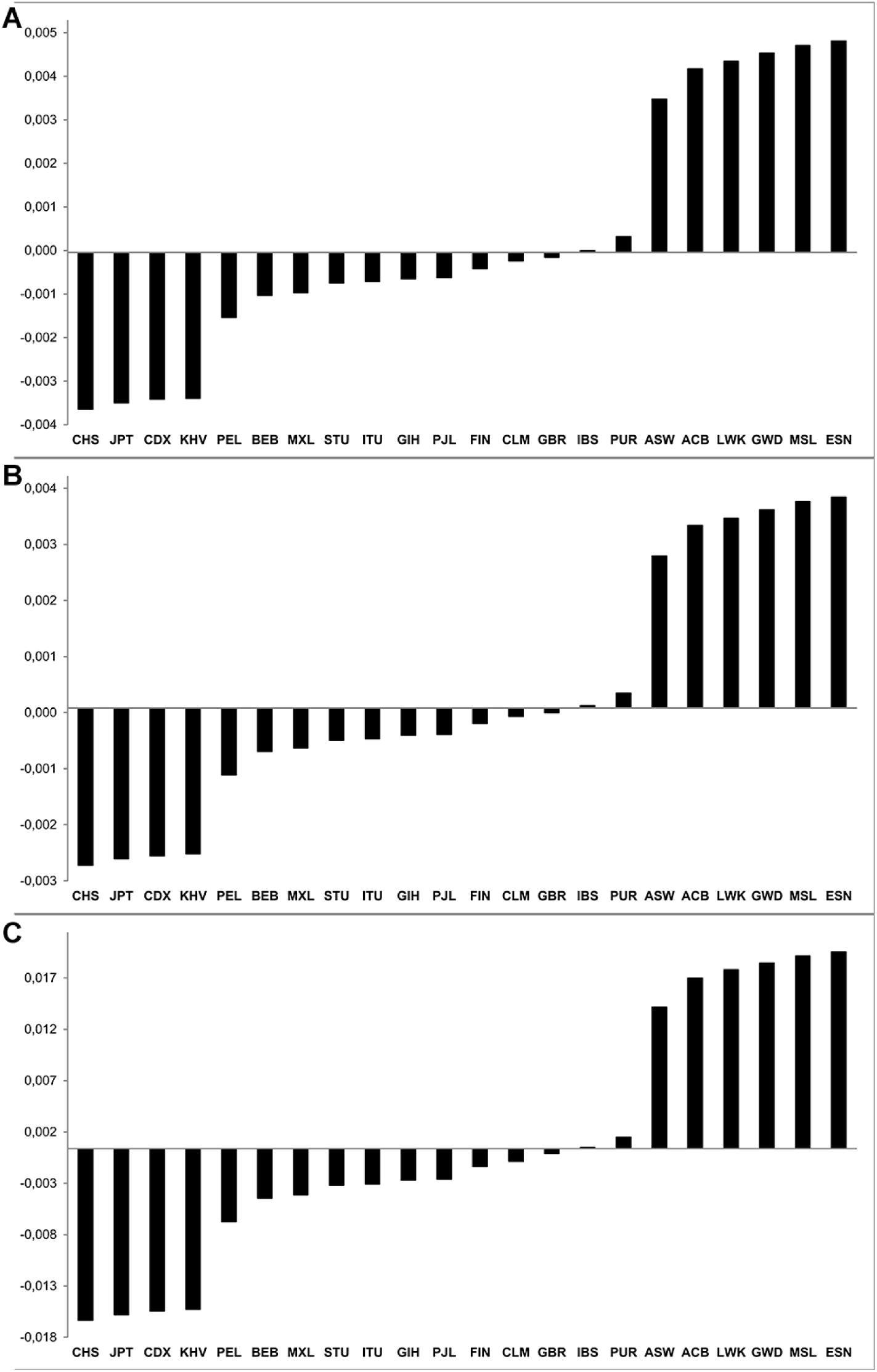
F_4_(TSI, X; CHB; YRI) values of the analyzed datasets: A) GENOME, B) EXOME and C) BEADCHIP, where X denotes the test population indicated on the x-axis.

In all analyzed populations, the *f*_*4*_-statistics showed nearly identical East Asian and African components for each dataset. A population with negative values indicates East Asian gene flow, while positive values indicate dominant African genetic components in the test population. The order of populations was the same for all datasets and the relative ratios between the F_4_ values of different populations was nearly identical. The absolute values were significantly higher in the BEADCHIP dataset compared to the GENOME dataset, while the EXOME dataset was more similar to it.

### Admixture analysis reveals subtle differences between the different datasets

We performed the admixture analysis and calculated the Cross Validation (CV) error for different models (K=3 to 10) for each dataset. Since the absolute CV error values were significantly higher in the BEADCHIP dataset compared to the GENOME and EXOME datasets, we displayed the relative CV values compared to the lowest fitting model (K=3) that resulted the highest CV error for each dataset (Figure 5).

**Figure 5:**
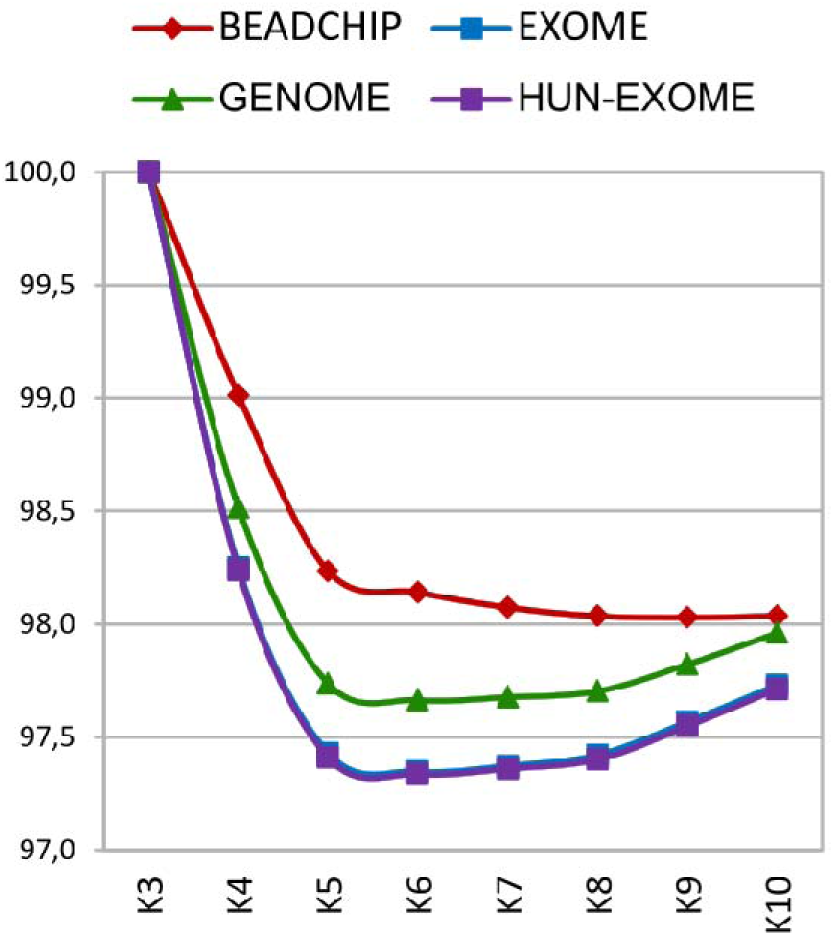
Calculated Cross Validation error for different admixture models (K=3 to 10) for each dataset.

The best fitting model of admixture indicated by the minimum of cross validation error was K=6 in the GENOME, EXOME and HUN-EXOME datasets, however the CV error of the BEADCHIP dataset indicated the K=9 model as the best fitting model of admixture. The curve of the CV errors of the BEADCHIP dataset shows that this dataset resulted in very similar alternative models (ranging from K=7 to 10) with almost identical CV errors. Since the analysis of the different dataset suggested different best fitting admixture models, we visualized both models (K=6 and K=9) for each dataset.

The analysis of the best fitting admixture model (K=6) suggested by both the GENOME and EXOME datasets produced very similar admixture results (Figure 6). Each color represents a different admixture component (K). The major topology and admixture components were very similar in each dataset for all analyzed populations however, the BEADCHIP systematically overestimated the minor admix components compared to the other two datasets (for example, South Asian component /marked as orange/ in Italian (TSI) or British (GBR) populations denoted by arrows in Figure 6).

**Figure 6:**
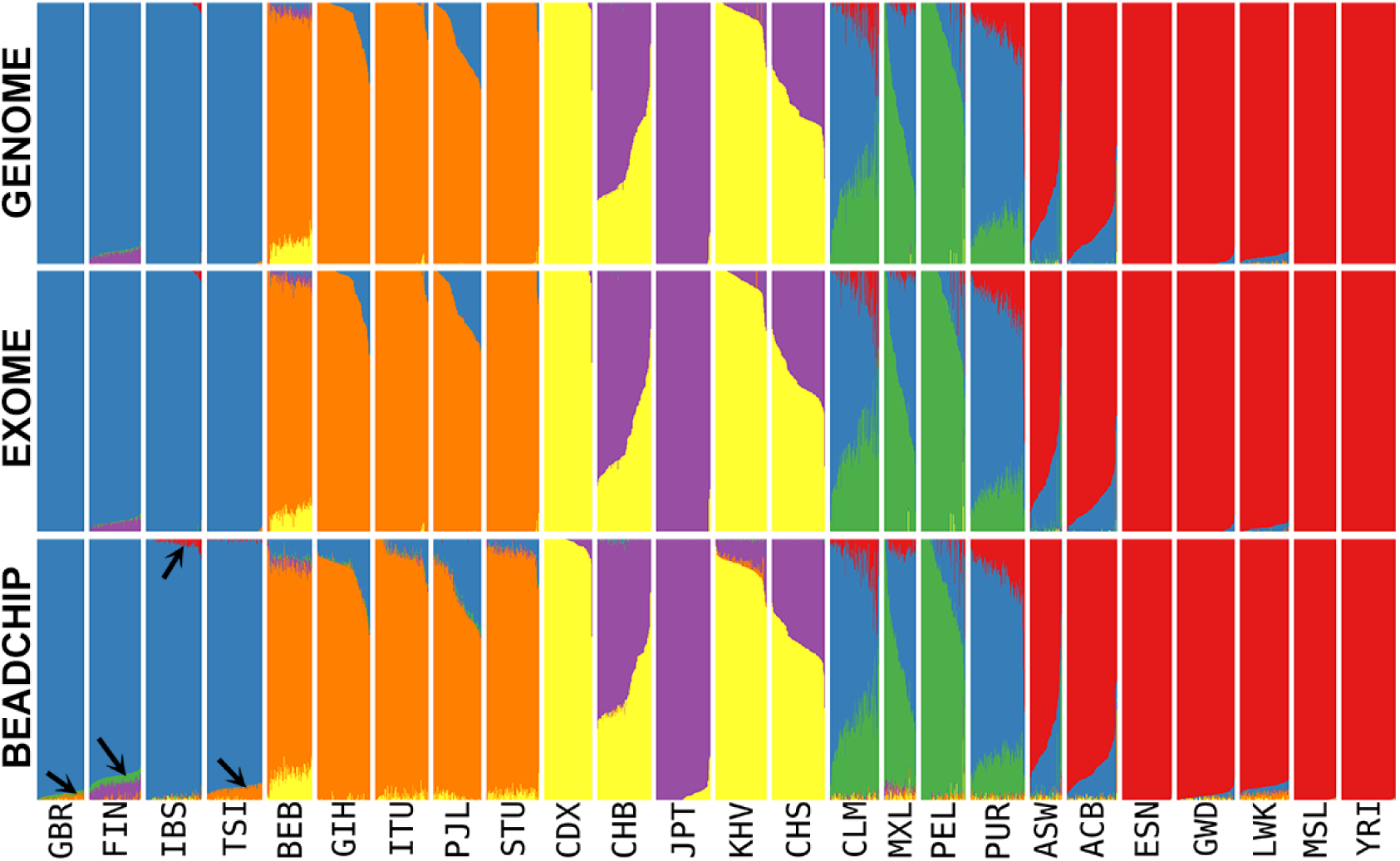
Admixture analysis of 25 populations using the GENOME, EXOME and BEADCHIP datasets with the K=9 admixture model. The arrows denote examples of minor genetic admixture components within the European super-population.

The analysis of the best fitting admixture model (K=9) suggested by the BEADCHIP dataset produced very similar admixture results for each datasets (Figure 7). The major topology and admixture components were very similar for each dataset. The major differences compared to the K=6 model were that in each dataset, the European super-population was split into two and the African super-population was split into three admixing components. The EXOME dataset was again most similar to the GENOME dataset, and again, the admixture of the BEADCHIP dataset systematically overestimated the minor admix components compared to the other two datasets (some examples are highlighted by arrows in Figure 7).

**Figure 7:**
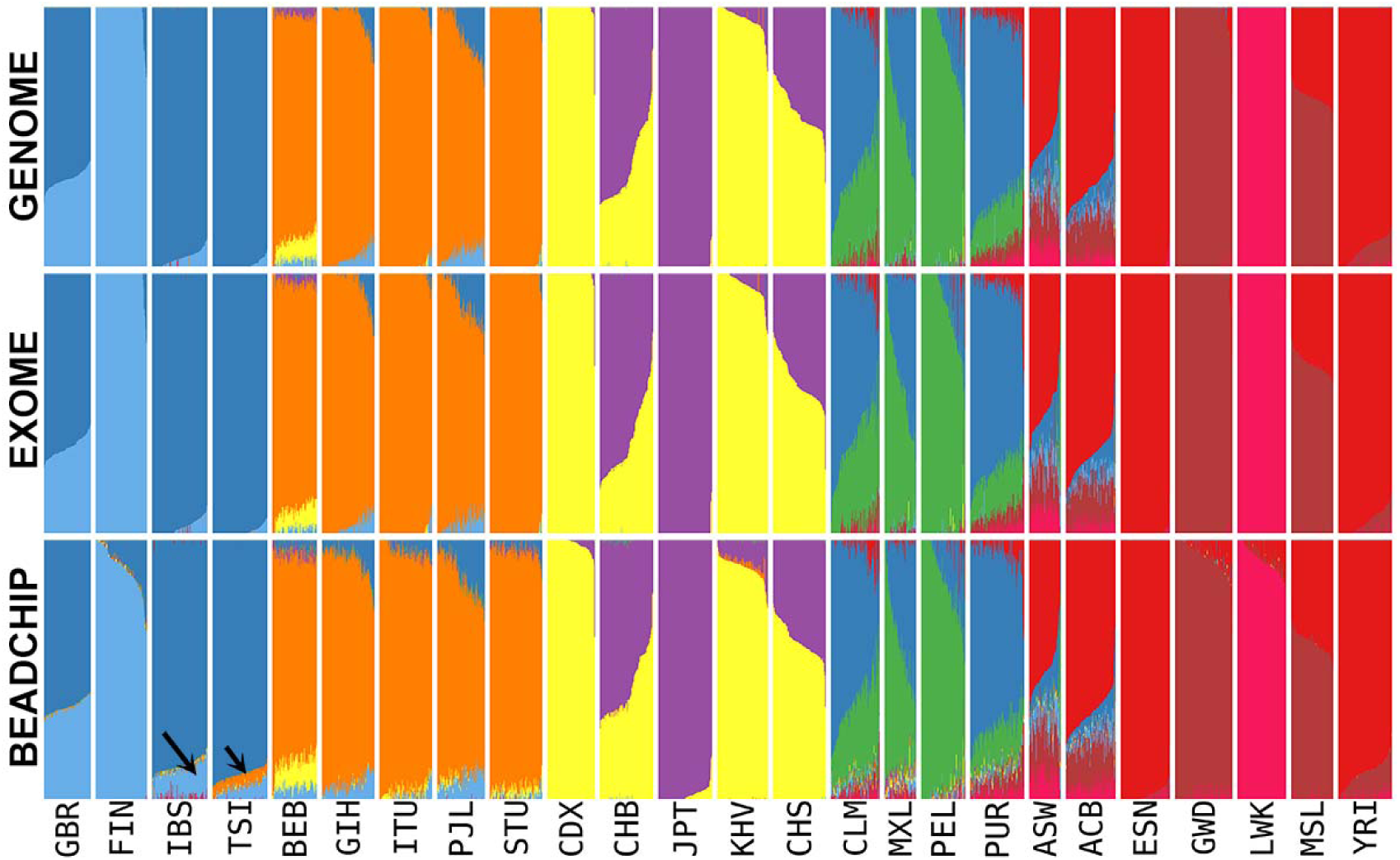
Admixture analysis of 25 populations using the GENOME, EXOME and BEADCHIP datasets with the K=9 admixture model. The arrows denote examples of minor genetic admix components within the European super-population.

### Overrepresentation of AIMs in the BEADCHIP dataset

Our previous population genetic analyses suggested that the variant composition of the BEADCHIP dataset is different from the GENOME and EXOME datasets. To test this hypothesis we calculated the variance of minor allele frequencies (MAF) of the analyzed populations for each SNP and visualized it as a density plot (Figure 8).

**Figure 8:**
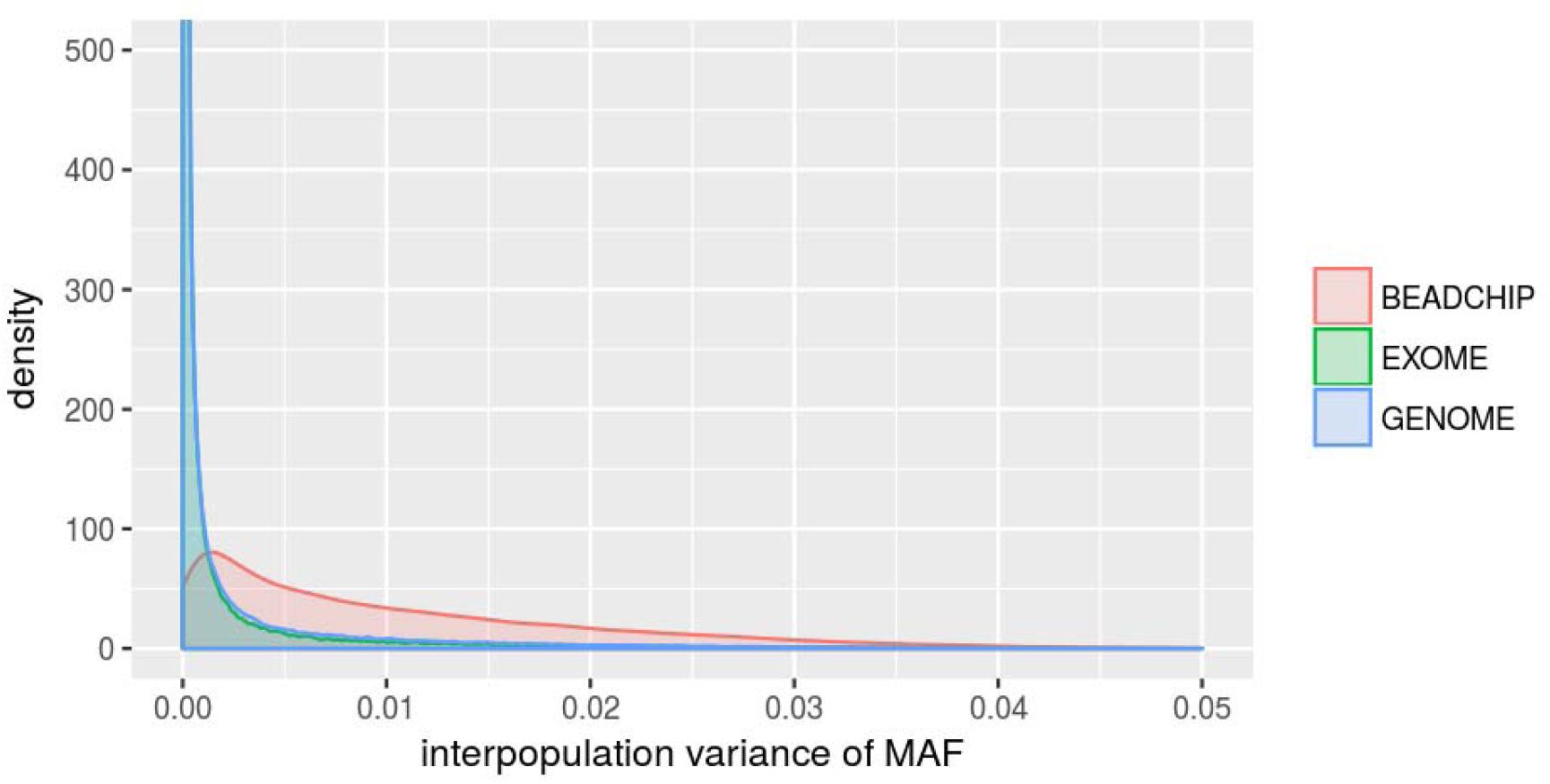
Density plot of the variation of the minor allele frequencies (MAF) of SNPs between the analyzed populations in the different datasets.

Figure 8 shows that SNPs that are highly variable between the test populations are overrepresented in the BEADCHIP dataset, while the distribution observed in the EXOME dataset is nearly identical to the distribution observed in the GENOME dataset. Thus, our analysis also confirms that EXOME dataset does not suffer from the same bias as the BEADCHIP dataset given correct processing of raw sequencing data for which we provide a step-by-step recommendation in Figure 9.

**Figure 9:**
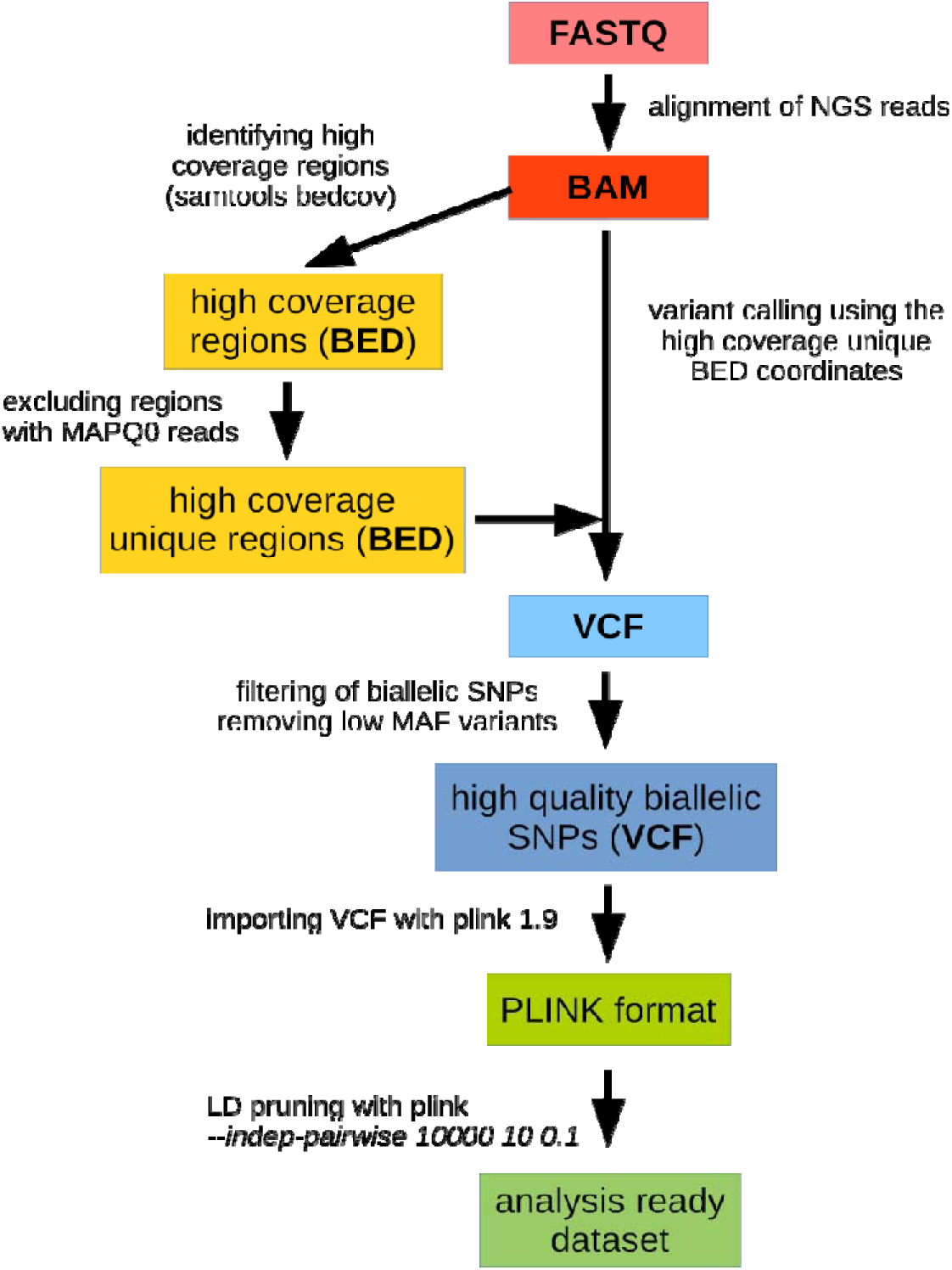
Preparation of WES data for population genetic analysis.

### Analysis of the HUN-EXOME dataset

The PCA of the HUN-EXOME dataset (Figure 10A) shows that Hungarians (denoted by purple circles) belong to the European super-population and that Hungarians are not in close relationship with the other European populations available in the public HGP dataset (Figure 10B).

**Figure 10:**
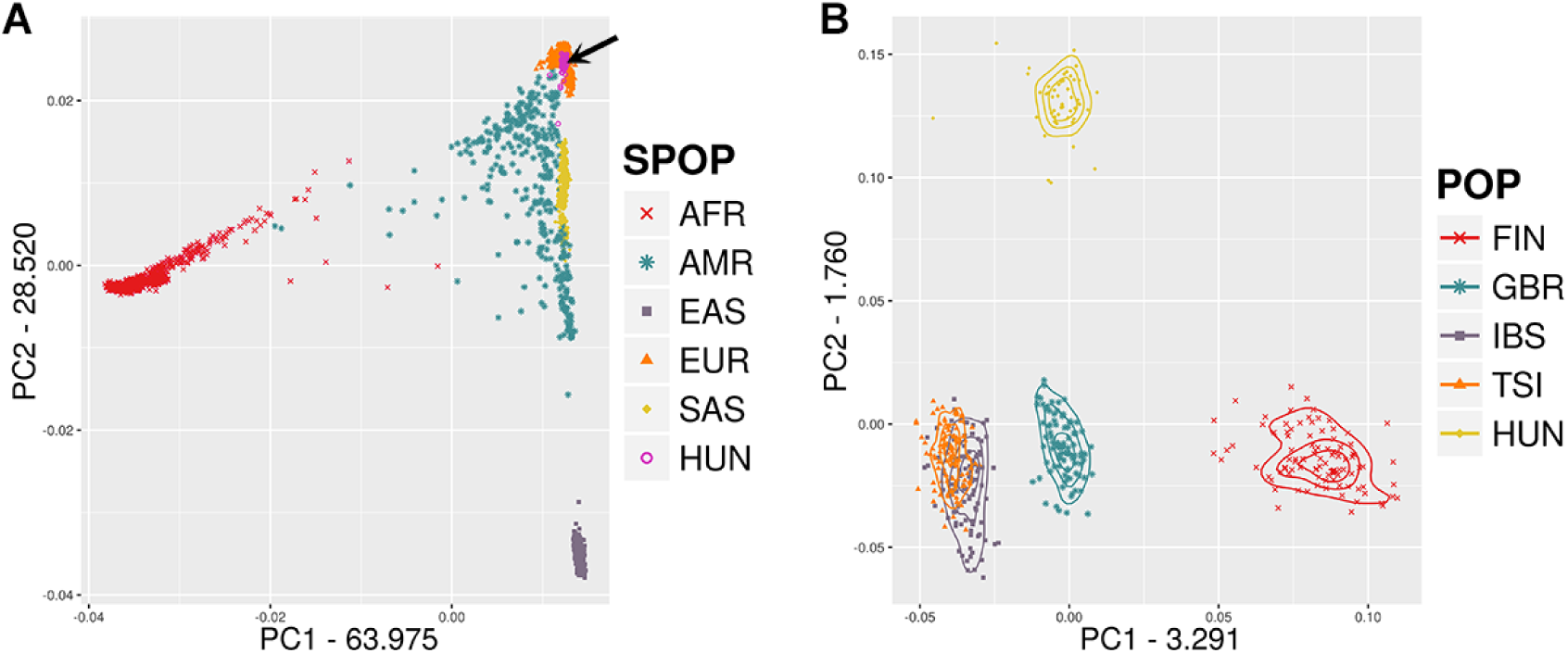
PCA of the HUN-EXOME dataset **A)** Hungarians (highlighted by the arrow) in relation to the super-population, **B)** Hungarians in relation to European populations.

In the case of Hungarians the F_4_ (TSI,HUN;CHB;YRI) value (Figure 11) showed that Hungarians have higher East Asian genetic components than the British population (GBR), but these genetic components are significantly smaller than those in the Finnish population (FIN).

**Figure 11:**
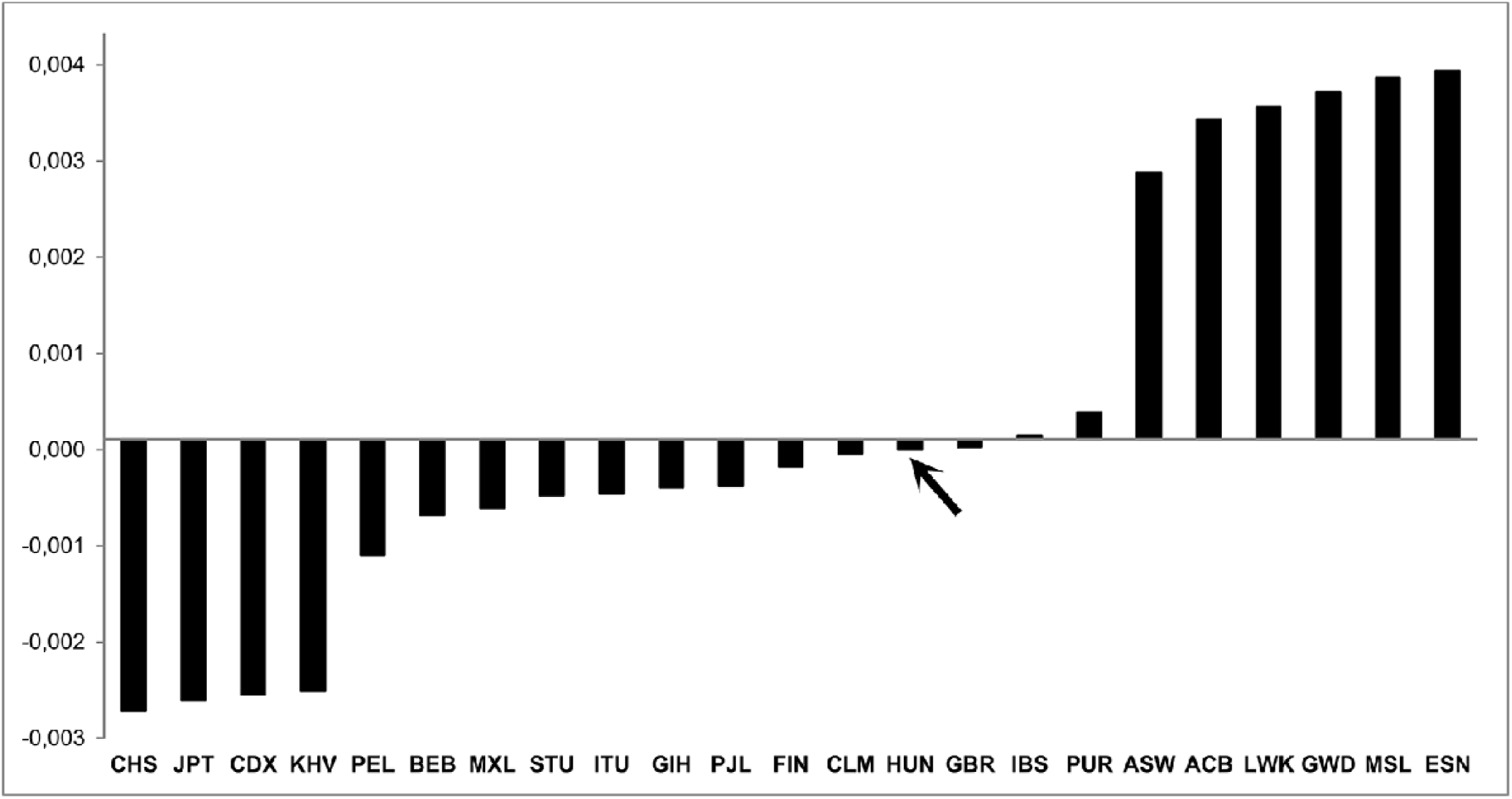
F4(TSI, X; CHB, YRI) values of the HUN-EXOME dataset, where X denotes the test population indicated on the x-axis. The arrow shows the relative position of Hungarians, indicating that they have higher East Asian admixture component than British but lower than Finnish population.

Since the analysis of the EXOME dataset was comparable to the other datasets, we also performed the admixture analysis of the HUN-EXOME dataset for both the K=6 and K=9 models.

In the K=6 model Hungarians were again classified into the European super-population as the major admixture component (depicted as blue) is the same as observed in other European populations (Figure 12A). Within the Hungarian population we can observe a few individuals with significant South Asian genetic components (denoted by orange). In the dataset of analyzed populations considering the K=9 admixture model (Figure 12B), Europeans display a North-South gradient by the indicated two European specific admix components (represented by the Finnish and Italian populations). According to their geolocation Hungarians are situated in the middle of this gradient having approximately 50-50% portion of these admix components.

**Figure 12:**
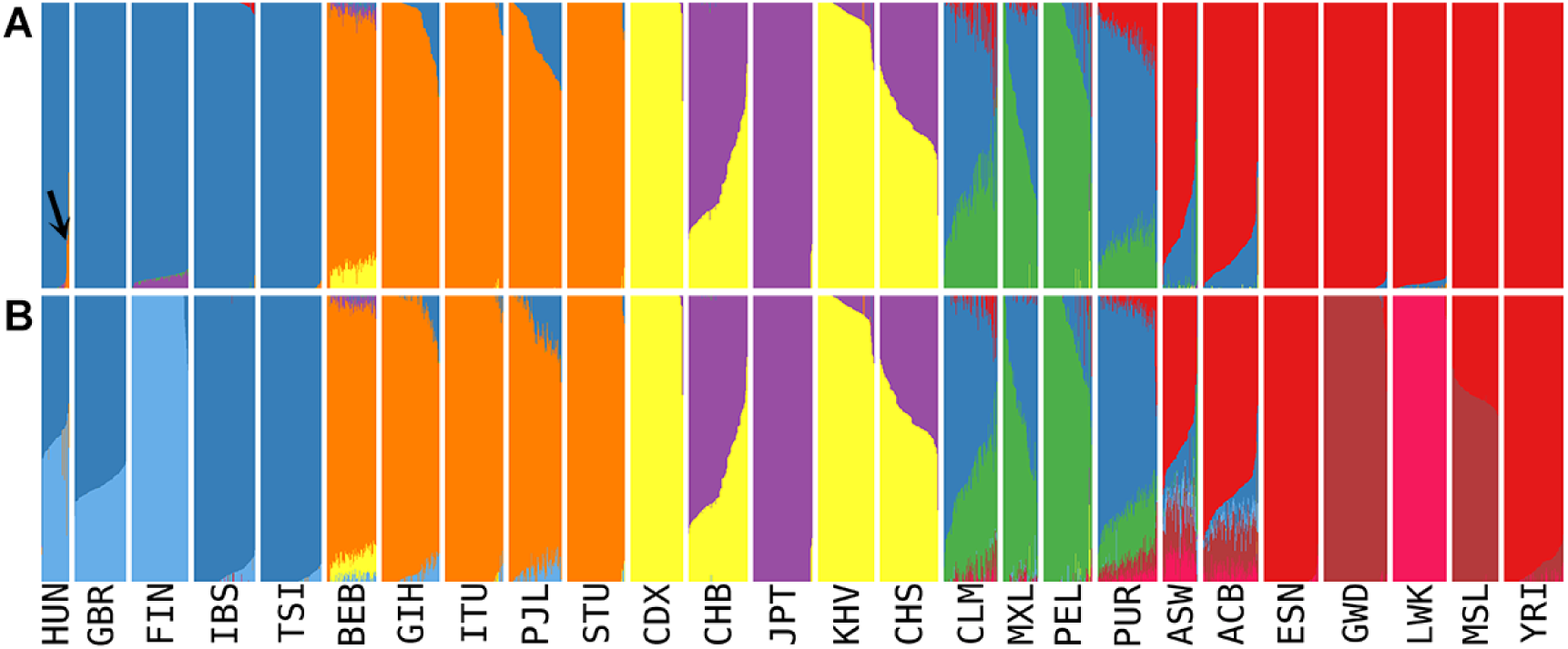
Admixture analysis of 25 populations using the HUN-EXOME dataset for the **A)** K=6 and **B)** K=9 admixture models. The arrow denotes Hungarian individuals with substantial South Asian admixture components.

## DISCUSSION

The WGS, the WES, and the array-based datasets are the three main types of human genetic datasets available today. In this study, we compared the use of these datasets for population genetic analysis, as each method differs in terms of the ratio in which it contains variants under natural selection. Our GENOME dataset mainly contains non-exonic variants, since more than 98% of the human genome consists of non-exonic region and our coordinate-based selection was random. The EXOME dataset contains both non-exonic (~50%) and exonic variants (~50%), although only a portion of the exonic variants are functional. In order to make the various approaches comparable, we used the same curated HGP 1kG genomic variant data to select the subsets of the GENOME, EXOME and BEADCHIP datasets (see in detail in the Methods section). The number of variants was comparable in all unpruned datasets (Table 1).

Proper LD pruning is a crucial step prior to PCA analysis, as large blocks of completely linked markers may introduce bias that could result additional eigenvectors (Patterson et al. 2006). Furthermore, it is also important in admixture analysis, because the calculations assume linkage equilibrium among the markers (Alexander et al. 2009). We observed that the unpruned BEADCHIP dataset contained slightly more variants than the GENOME dataset (567k vs. 483k variants), but most of them were tightly linked, as only ~72k markers (~12%) remained after LD pruning, while in the GENOME dataset the ratio was ~43% (~205k markers) which indicates a smaller fraction of linkage. We suppose that these differences are contributed to the tightly linked pre-selected AIMs in the BEADCHIP dataset. The coordinate-based EXOME dataset had somewhat higher linkage than the dispersed GENOME dataset. This is assumed to be a consequence of the organization of the human genome, where genes and exons are not homogeneously dispersed, but rather tend to be packed tightly in functionally active euchromatic chromosome regions. Correspondingly, after LD pruning about 149k, a slightly lower proportion of variants (~38%) remained in the EXOME dataset out of the ~405k unpruned variants. Comparing the PCA results of the EXOME dataset with the gold standard GENOME dataset, we refined the LD pruning parameters of exome data. We suggest extending the pruning window (to 10,000 kb) – while keeping the original 0.1 squared correlation threshold – in order to counter the effect of the packed exome variant composition and to eliminate the tightly linked markers. According to our results, this modification did not significantly alter the variants of the BEADCHIP dataset; however, it did eliminate additional tightly linked variants in the EXOME and to a lesser degree in the GENOME datasets (Table 1).

The fixation index had been developed as a special case of Wright’s F-statistics and is one of the most commonly used statistics in population genetics, which is a measure of population differentiation due to genetic structure. F_ST_ calculates the overall reduction in average heterozygosity between populations. F_ST_ calculations of the GENOME and EXOME datasets resulted in nearly identical pairwise genetic distances between the analyzed populations. However, we found that the F_ST_ distances between African (except the LWK population) and the European populations were very slightly (0.001-0.002) but systematically smaller using the EXOME dataset. On the other hand, the F_ST_ values between the African and East Asian populations were very slightly (0.001 - 0.002) but systematically larger compared to the GENOME F_ST_ distances. Since this slight difference was systematic between the random GENOME and the EXOME dataset (which by definition contains functional variants besides the non-coding and other functionally inert variants), we hypothesize that a portion of functional variants are accountable for this phenomenon. However, the deviation (~1-3%) is still only a portion of what was observed in the BEADCHIP dataset. The comparison of F_ST_ values of the BEADCHIP dataset to the GENOME dataset revealed that the pairwise F_ST_ distances between populations of different super-populations were systematically larger (~1-12%), and that the extent of the difference appears to be correlated to the phylogenetic distance. On the other hand, we detected almost no differences between the F_ST_ distances of populations within the same super-population, except in the highly admixed AMR super-population. We assume that this is due to a general overrepresentation of differentiating SNPs (AIMS) and imbalances in the selection and proportion of the marker composition in the pre-selected BEADCHIP dataset. This hypothesis is also supported by the observed F_ST_ values of the admixed ASW population. The ASW population is a sub-population of African Americans in the Southwestern United States who originated from West-Africa, and later mixed with Caucasian and American Indians. Accordingly, the BEADCHIP data places this population closer to its admix sources - the European (EUR) and Admixed American (AMR) super-populations - than those of the GENOME F_ST_ values, indicating that the overrepresentation of AIMs distorts the true population distances. As F_ST_ can be used to estimate coalescence times (Weir and Cockerham 1984; Hudson et al. 1992), significant deviations in F_ST_ values - such as observed using the BEADCHIP dataset - may lead to bias in these estimations.

For each dataset the PCA analysis showed both similar topology and comparable relations between the analyzed populations. We observed that for the first two eigenvectors of the whole dataset (Figure 1A-the eigenvalues were greater in the BEADCHIP dataset compared to the GENOME dataset, while the EXOME datasets were more similar to it. The eigenvalue with the largest absolute value is known as the dominant eigenvalue and can be used to determine the rate of growth in the population (Caswell 2000). An overestimation of the eigenvalues miscalculates the population growth rate. The largest differences between the detailed PCA were seen in the African super-population (Figure 2E) where the BEADCHIP dataset gave slightly different eigenvalues and population relations compared to the GENOME and EXOME datasets. Similarly to the F_ST_ results, the admixed ASW and ACB populations displayed slightly different relations to other populations. We assume that again, the overrepresentation of AIMs in the BEADCHIP dataset is responsible for the increased eigenvalues and the different relations of the admixed ASW and ACB populations. Nonetheless, the genetic relationship among the studied populations was still comparable.

The 3-population test is a formal test of admixture and can provide clear evidence for admixture, even if the gene flow events occurred hundreds of generations ago (Patterson et al. 2012). Positive *f*_*3*_-statistics are consistent with either a simple tree or a history of admixture followed by genetic drift, while significantly negative *f*_*3*_-statistics reveal that the target and the two parent populations do not form a simple tree, indicating complex relationships. In order to test the usability of each dataset for all of the potential relationships between the populations indicated by the *f*_*3*_-statistics (tree like, admixture, or no relation), we generated all population triads and their corresponding *f*_*3*_-statistics for each dataset. Comparison of the three datasets showed that the GENOME and the EXOME data gave almost identical F_3_ values (r=0.9998), while the Z scores were slightly smaller in the EXOME dataset, which was attributable to the smaller SNP count. On the other hand, the *f*_*3*_-statistics of the BEADCHIP dataset showed less correlation (r=0.9911). The Z scores of the *f*_*3*_-statistics in the BEADCHIP dataset were also higher, even though the SNP count was about one third of the GENOME dataset, which indicates higher deviation from the mean. Plotting the corresponding F_3_ values revealed systematic differences in a large portion of population combinations between the BEADCHIP and GENOME datasets. Investigation of F_3_ values showed that tree like split of East Asian populations from African populations was systematically overestimated by the *f*_*3*_-statistics based on the BEADCHIP dataset. We assume that this bias is due to the overrepresentation of AIMs between the East Asian and African populations within the pre-selected variants of the BEADCHIP dataset. Taking all this together, we conclude that the EXOME data is highly suitable for *f*_*3*_-statistics. Although the absolute F_3_ value deviation from zero is meaningless if we only test whether *f*-statistics is consistent with zero (Patterson et al. 2012) (meaning no admixture), the inferred magnitude of relatedness in comparison to relations between other populations may lead to bias or overinterpretation of data in some specific cases (such as where markers are not evenly represented in the analyzed populations) of the BEADCHIP dataset (due to higher F_3_ values and optimistic Z scores).

In the case of the *f*_*4*_-statistics, we observed nearly identical relative orders and ratios in the analyzed sample of East Eurasian ancestry in Northern European populations between the three datasets. The only difference was the absolute F_4_ values, in which case the BEADCHIP dataset resulted in higher F_4_ values, while the EXOME dataset derived values similar to those of the GENOME dataset. Since *f*_*4*_-statistics is used for the estimation of the admixture proportions of a test population, the absolute F_4_ values themselves are meaningless. Our analysis also supports that *f*_*4*_-statistics is robust to SNP ascertainment (all three datasets resulted in nearly identical proportions) and deviation from zero is only observed if the test population is admixed (Patterson et al. 2012).

The admixture analysis of different datasets suggest different best-fit admix models. The cross validation (CV) error was lowest and thus, the suggested best fitting admixture model was K=6 in the GENOME and the EXOME datasets. In the BEADCHIP dataset, the best fitting model with the smallest CV error was K=9. Additionally, for this dataset, the K=7-10 models had almost identical CV errors (Figure 5). Since the BEADCHIP dataset resulted in higher CV errors and a number of very similar alternative models, it appears that this dataset is less conclusive for pinpointing the true admixture model. Nevertheless, the admixture analysis of the different datasets resulted very similar admixture components for all of the tested models (Figure 6-7). We observed that the ratio of minor admixture components were overrepresented in the admixture analysis of the BEADCHIP dataset compared to the GENOME dataset. On the other hand, the admixture results of the EXOME dataset were very similar to the GENOME dataset, although in some cases the minor admixture components were slightly underrepresented.

In our comparative analyses, we used the same HGP dataset to make an unbiased assessment of the different strategies of variant selection. However, comparing datasets from different sources may lead to bias due to the differences in the applied NGS variant calling tools, pipelines, thresholds, and quality of sequences. The majority of genotyping uncertainties stem primarily from different quality parameters and thresholds applied to call variants in the various NGS pipelines and the changing information content of certain genomic regions (due to inter-sample variances caused by target enrichment). Thus, in order to minimize the bias, we applied a number of criteria for exome data processing before using them for population genetic analyses. Hence, we excluded the low-coverage data, based on the real coverage profile of our aligned NGS reads, as these data may have led to ambiguous variant calls. We excluded all read length variations (INS/DELS/DUPS) from the analysis since the same variant may be represented ambiguously by different variant callers. We also excluded pseudogenic regions where the proportion of the alternatively aligned sequencing reads may lead to differences depending on the threshold values or the pipeline applied for the analysis.

Using these strict criteria, we merged the Hungarian exome data and analyzed the resulting HUN-EXOME dataset with the appropriate population genetic tools. Both the PCA and admixture analysis of the modern Hungarian exome dataset confirms that genetically, modern Hungarians are Europeans (Figure 10 PCA; Figure 12 K=6, K=9 admixture plots). We note here that our admixture analysis (K=9) indicated two genetic components with a North-South European gradient which has a ~ 50-50% portion in Hungarians. Admixture analysis also detected a portion of a South Asian component in a few individuals of the Hungarian population, which in our view can be attributed to the Gypsies living as an ethnic minority (~5%) in Hungary. F_3_ analysis of HUN-EXOME dataset also suggests a small but significant (high Z scores) South-Asian-HUN admixture (Table 2).

**Table 2:** *f*_*3*_-statistics of Hungarian and selected South Asian populations.

| Pop A | Pop B | Pop C | F3 value | Std Err | Z score | SNP count |
| --- | --- | --- | --- | --- | --- | --- |
| STU | HUN | PJL | -0.003761 | 0.000140 | -26.782 | 76834 |
| ITU | HUN | PJL | -0.003641 | 0.000141 | -25.752 | 76845 |
| BEB | HUN | PJL | -0.002865 | 0.000138 | -20.743 | 80622 |

Negative $F_3$ values with significant Z score indicate complex relations (admixture) between the Hungarian and South Asian populations.

Historically, gypsy tribes left India in the 9^th^ and 10^th^ centuries as a result of Muslim attacks in areas they inhabited and first appeared in territory of the medieval Kingdom of Hungary in the 14^th^ and 15^th^ centuries, probably fleeing from the conquering Turks in the Balkans (Kemeny). This assumption is also supported by the fact that the South-Asian genetic component is confined to only a few Hungarian individuals with high contributions indicating a recent admixture. Unfortunately we cannot explicitly verify the ethnic origin as our exome data was anonymous and therefore we had no information on ethnicity. Modern Hungarians identify themselves as having originated from the Hungarian Conquerors, who are deemed to have arrived to the Carpathian Basin in the 9^th^ Century migrating from around the Ural Mountains, which is the natural border between Europe and Asia. The *f*_*4*_-statistic indicated a small East Asian admixture component in modern Hungarians (Figure 11) that was also seen as minor admixture components in few Hungarian individuals (Figure 12). The minor East Asian component detected in modern Hungarians is possibly the genetic trace of Hungarian Conquerors as also suggested by mtDNA analysis (Tomory et al. 2007; Biro et al. 2015). Because the publicly available HGP dataset contains worldwide populations with no or very little genetic relation to modern Hungarians, our analysis indicated only known and plausible relations of modern Hungarians in relation to the analyzed populations (such as general European ancestry with small-scale East-Asian components and a recent admixture with South-Asian components). However, this systematic analysis confirms the usability of WES data in population genetic analysis and leads us to conclude that exome data from populations having shared ethnogenesis with Hungarians could result in an even better understanding of their genetic history.

Overall, our comparative analysis indicates that the array-based preselected SNP-set (BEADCHIP dataset) deviates most from the GENOME dataset. The reason behind this phenomenon may be twofold. First, the pre-selection of SNPs in the array is not necessarily a uniform representation of all of the analyzed populations, and second, the increased proportion of AIMs is shifting the balance towards highlighting the differences between populations. The increased proportion of AIMs makes the BEADCHIP dataset suitable for sensitive exploration of admixture components. However, it could also lead to deviations of the Fixation index, specific cases of f3-statistics, PCA eigenvalues, suggested admixture model, and admixture rates. Thus, all derived parameters (e.g. coalescent time, population growth rate) estimated from these statistics are prone to bias in case of array-based data. WES data, on the other hand, is not affected by this bias and is suitable for population genetic analyses. Based on the usability of the EXOME dataset we would encourage everyone to use and to publicly share WES sequences with the correct indication of ethnic and geographical origin, which could contribute towards a better understanding of the genetic relationships among human populations.

## METHODS

### Preparation of Hungarian exome data

Our previously published BAM files of Hungarian whole exomes (Tombacz et al. 2017) were aligned by the BWA-MEM algorithm (Li 2013). Using the whole cohort, we identified the high coverage regions (all samples in the cohort had coverage >10x) using *samtools bedcov* (version 1.3.1) algorithm (Li et al. 2009) with the SureSelect V5 all exon plus UTR kit manifest bed coordinates. Since some of the regions may contain repetitive elements or pseudogenic regions with non-unique sequences, we excluded all regions that had any *MAPQ0* reads (*mapping quality 0*, indicating that the read could be mapped to multiple genomic regions). We excluded sex chromosomes from the analyzed regions. As a result, we generated a BED coordinate list (Supplementary Data/HighCov_HighQual_EXOME.bed) that contained the high coverage, unique genomic regions of the exome kit. Variant calling was performed by the *GATK HaplotypeCaller* (version 3.5) best practice (Van der Auwera et al. 2013) using the parameters *-stand_emit_conf 10 -stand_call_conf 30* in the HaplotypeCaller module, and *-- ts_filter_level 99.0* in the variant recalibration (ApplyRecalibration) module. We filtered the variants to include only high quality SNPs (PASS filter) in the final high coverage/unique bed regions. We also filtered out multi-allelic variants and variants that had < -0.5 *InbreedingCoeff* (indicating variants that violate HW equilibrium, which would potentially be technical errors since our cohort consisted of unrelated patients).

### Preparation of public datasets

We used the publicly available variants of the Human Genome Project 1kG phase 3 dataset (Genomes Project et al. 2015). We excluded the geographically inaccurate CEU population and all of the related individuals from our analysis. Our final dataset included sequence data of 2,369 individuals originating from 25 available populations, as follows: European (EUR: GBR - British in England and Scotland; FIN - Finnish in Finland; IBS - Iberian populations in Spain; TSI - Toscani in Italy), South Asian (SAS: BEB - Bengali in Bangladesh; GIH - Gujarati Indian in Houston TX; ITU - Indian Telugu in the UK; PJL - Punjabi in Lahore, Pakistan; STU - Sri Lankan Tamil in the UK), East Asian (EAS: CDX - Chinese Dai in Xishuangbanna, China; CHB - Han Chinese in Bejing, China; JPT - Japanese in Tokyo, Japan, KHV - Kinh in Ho Chi Minh City, Vietnam; CHS - Southern Han Chinese, China), Admixed Americans (AMR: CLM - Colombian in Medellin, Colombia; MXL - Mexican Ancestry in Los Angeles, California; PEL - Peruvian in Lima, Peru; PUR - Puerto Rican in Puerto Rico), and African (AFR: ASW - African Ancestry in Southwest US; ACB - African Caribbean in Barbados; ESN - Esan in Nigeria; GWD - Gambian in Western Division, The Gambia; LWK - Luhya in Webuye, Kenya; MSL - Mende in Sierra Leone; YRI - Yoruba in Ibadan, Nigeria).

To compare the capabilities of the sequencing-based (WES and WGS) and array-based approaches, we created three SNP datasets (denoted as EXOME, GENOME and BEADCHIP) established upon the following rules: first, the EXOME dataset (Supplementary Data/EXOME) was prepared by filtering the public 1kG variants with the *bedtools intersect* algorithm using the genomic coordinates of the high-coverage high-quality exome BED coordinate list that we established using the Hungarian exome data. Second, the GENOME dataset was created on the basis of a homogeneously dispersed genomic coordinate list that spanned all somatic chromosomes (Supplementary Data/GENOME.bed). The first and last 5Mb of the telomeric regions of chromosomes were excluded in order to eliminate uncertain sequences, such as repetitive elements. The remaining genomic regions on each chromosome was equally divided; the first 258 base pairs in every 10kb DNA stretch were used by the *bedtools intersect* algorithm for extracting homogeneously dispersed variants across the genome (Supplementary Data/GENOME). The resulting unbiased genomic region coordinates were found to have a cumulative size (68.7 Mb), which is similar to the cumulative size of the high-coverage exome regions (68.9 Mb).

In order to prepare the BEADCHIP dataset (Supplementary Data/BEADCHIP), the genome coordinates of a frequently used Human genotyping array kit (Illumina Human610-Quad BeadChip) were used from the kit’s manifest file and converted to BED coordinates. The *bedtools intersect* algorithm was used to filter the corresponding variants from the 1kG genomic variants. From all three datasets, we removed all multi allelic variants, along with variants that were found in less than 10 individuals (or in all but 10 individuals in case of refseq error). This resulted in comparable variant numbers for each dataset (567k BEADCHIP, 405k EXOME, 483k GENOME). The 1kG annotated joint VCF was re-coded to *plink.ped* and *plink.map* formats. Plink (version 1.9) was used to generate the binary bed format files used in the downstream analysis (Chang et al. 2015).

### Merging Hungarian exome data with public dataset

In order to create the merged HUN-EXOME dataset, we used the same high-coverage high-quality exome BED coordinate list, which we established using Hungarian exome data, and the *bedtools intersect* algorithm for filtering the variants in both the Hungarian exome data and the 1kG datasets. The filtered exome and 1kG variants were merged with the *GATK CombineVariants* algorithm. We removed multi allelic variants, as well as variants that were only found in less than 10 individuals (or all but 10 individuals in case of refseq error). The resulting joint VCF were re-coded to *plink.ped* and *plink.map* formats and we used plink to generate the binary bed format file (Supplementary Data/HUN-EXOME).

### LD pruning of datasets

LD pruning was carried out using the *–indep-pairwise* algorithm of plink (version 1.9). Because of the structure of the human genome, it was expected that exome variants would yield non-homogeneously dispersed markers. Therefore, we performed the LD pruning using different sliding window sizes (the recommended 50kb of admixture protocol, and an extended 10,000kb) to prune linked markers in order to identify the best parameters. For both window sizes, we kept the recommended 10 SNPs increment and r^2^ threshold of 0.1 to allow pruning of markers by the same linkage criteria. In order to allow unbiased comparison of the three methods, we used the same LD pruning parameters (10,000kb sliding window, 0 SNPs increment and r^2^ threshold of 0.1) for each dataset.

### *F*_*ST*_ and PCA calculations

The pairwise F_ST_ matrix and the PCA analysis of populations were performed by the EIGENSOFT software (version 6.1.3) (Patterson et al. 2006) on the LD pruned datasets. For each subset of data (whole dataset and the different super-populations), the LD pruning of linked variants and the subsequent PCA calculations were performed separately. PCA analysis was carried out without outlier removal for the whole datasets and with outlier removal (SD=6) for the analysis of individual populations for each super-population. PCA plots were visualized by ggplot2 (version 2.2.1) (Wickham 2009).

### *f*_3_ and *f*_4_-statistics

The F_3_ tests were carried out by the *qp3Pop* program of ADMIXTOOLS (Patterson et al. 2012) for each population triad of the analyzed populations. The F_4_ tests were calculated using the *fourpop* algorithm of TreeMix software (version 1.13). For each dataset, the corresponding F_4_(TSI,X;CHB,YRI) values were calculated (where X refers to the given test population). F_3_ and F_4_ values of the tests were visualized by ggplot2.

### Admixture analysis

Admixture analysis was performed by the ADMIXTURE software (version 1.3.0) using the LD pruned datasets with the *--cv* option for K=3 to 11 values using 20 iterations and randomized seeds (Alexander et al. 2009). The best admixtures suggested by the cross-validation plots of genome/exome and BEADCHIP datasets were visualized by a custom Perl script using Linux ImageMagick software (version 6.7.7-10).

